# Virophage replication mode determines ecological and evolutionary changes in a host-virus-virophage system

**DOI:** 10.1101/2024.04.15.589535

**Authors:** Ana del Arco, Lutz Becks

**Affiliations:** Aquatic Ecology and Evolution Group, Limnological Institute, University of Konstanz, Germany

## Abstract

Giant viruses can control their eukaryotic host populations, shaping the ecology and evolution of aquatic microbial communities. Understanding the impact of the viruses’ own parasites, the virophages, on their control of microbial communities remains a challenge. Most virophages have two modes of host infection and replication. They can exist as free particles that co-infect a host cell with the virus and replicate but inhibit viral replication. Virophages can also integrate into the host genome and remain dormant until the host is infected with a virus, leading to virophage reactivation and replication that does not immediately inhibit viral replication. Both replication modes are present within host-virus-virophage communities, and their relative contributions are expected to be context dependent and dynamic over time. The consequences of this dynamic regime for ecological and evolutionary dynamics remain unexplored. Here, we test whether and how the relative contribution of virophage replication modes influences the ecological dynamics of an experimental host-virus-virophage system and the evolutionary responses of the virophage. To do this, we indirectly manipulated the level of virophage (Mavirus) integration into the host (*Cafeteria burkhardae*) in the presence of the giant Cafeteria roenbergensis virus (CroV) (later identified as *Cafeteria burkhardae*, Schoenle et al. 2020). Our results show that higher virophage integration is positively correlated with host survival, but negatively correlated with virophage reactivation. In addition, communities with higher virophage integration were characterised by lower population densities and reduced fluctuations in both host and viral populations, whereas virophage fluctuations were increased. This study reveals the complex interplay between virophages, viruses and hosts, in which the virophage dual replication mode is a dynamic and reactive mechanism contributing to persistence of the microbial community.

## Introduction

Aquatic viruses play a central role in marine microbial community dynamics, biogeochemical cycling, and diversification (Breitbart et al. 2007; Suttle 2007; Biggs et al. 2021). Understanding how the role of viruses is influenced by their own parasites, the virophages, remains a challenge (Duponchel and Fischer 2019). Many virophages can adopt a dual lifestyle in which they either integrate into the genome of the host of the virus, or they exist as free particles that can co-infect the host together with the virus (Berjón-Otero et al. 2019). Integrated virophages are transmitting vertically through the replication of the host. They can be reactivated upon viral infection of the host and parasitize the virus by using its virus factory for its own replication and production of free virophage particles (La Scola et al. 2008; Fischer and Hackl 2016). During co-infection, the virophage also exploits the virus factory of the virus for its own replication, allowing for horizontal transmission. While the importance of virophages for host and virus dynamics has been demonstrated in a few studies (Wodarz 2013; Taylor et al. 2014; Yau et al. 2016; Fischer and Hackl 2016; del Arco et al. 2024), the ecological consequences of this viral exploitation by the virophage for the microbial community remain largely unknown.

The mode of virophage replication can be an important driver of ecological and evolutionary responses of host-virus-virophage dynamics. For example, coinfections in which virophages enter the host cell independently, as in the case of the Mavirus virophage (Fischer and Hackl 2016), or in which they enter together with the virus, as in the case of the sputnik virophage (La Scola et al. 2008), lead to different population dynamics (Taylor et al. 2014). When virophages can transmit vertically through host cell division or horizontally from one infected host cell to other host cells, as in the case of Mavirus virophages (Berjón-Otero et al. 2019), the mode of virophage replication affects virus replication: during reactivation of integrated virophages, virus replication is unaffected compared to infections in the absence of virophages, whereas during co-infections, virus replication is reduced. These differences in virus replication have implications for population dynamics, for example by driving virus densities to very low levels (Fischer and Hackl 2016).

In the absence of an infectious virus, the virophage’s mode of replication is fixed as vertical. In addition to vertical transmission with host replication, the virophage has two replication modes - reactivation of integrated virophage and co-infection – and they are expected to be present in a community if infectious virus is present. Their relative contributions may change depending on population densities, traits that determine the outcome of the interaction, and environmental conditions. Host, virus and virophage densities change over time affecting encounter rates between hosts and viruses and thus the likelihood for co-infections. In addition, traits underlying virus and virophage replication and host exploitation may evolve rapidly. For example, previous work with the Cafeteria-CroV-Mavirus system has shown that the high degree of exploitation and interdependence in this tripartite system leads to the rapid evolution of reduced replication of the virus (CroV) and increased virophage (Mavirus) replication, but reduced exploitation of the virus by the virophage (del Arco et al. 2024). These observations together suggest that the virophage replication mode is not constant over time and that the relative contribution of the two modes to virophage and virus reproduction may continuously change. The consequences of this dynamic regime for ecological and evolutionary dynamics are unexplored.

As a first step for developing an understanding of these consequences we test here the hypothesis that differences in the frequency of integrated virophage determine ecological (population dynamics) and evolutionary dynamics (virophage replication and viral exploitation by the virophage) through their effect on virus replication. We predict higher virus and virophage densities in communities with higher levels of virophage integration because both viruses can replicate after reactivation of integrated virophages, which also leads to lower host densities. The long-term persistence of such a tripartite system should favor virophage traits that allow adaptations for sufficient exploitation of the virus without compromising the abundance of its viral host as a resource (Morozov et al. 2007; Heilmann et al. 2010). This can be achieved if the virophage evolves higher levels of replication and lower levels of exploitation of the virus (del Arco et al. 2024), as high levels of exploitation would result in low virus densities over time and thus few opportunities for the virophage to parasitize the virus. With more integrated virophages per host, we predict that selection for reduced exploitation will be weaker compared to hosts with fewer integrated virophages. This is because reactivation of integrated virophages leads to virus production and thus, on average, higher virus densities, whereas co-infection only leads to virophage production and thus lower virus densities. Since previous work has shown that increased reproduction and reduced virus exploitation by the virophage evolve together (del Arco et al. 2024), we further predict that virophage reproduction will either not evolve or evolve with a smaller increase.

We tested the hypothesis in experiments with the host *Cafeteria burkhardae*, the giant virus Cafeteria roenbergensis virus (CroV) and the virophage Mavirus and manipulated the level of integration of the virophage into the host genome, by adding a chemical stressor to the experimental communities. Preliminary observations showed higher levels of virophage integration into the host genome resulting from an off-target effect of the antiviral oseltamivir (del Arco *personal observation*). We used the addition of the antiviral as an approach to manipulate the level of virophage integration rather than different host strains with naturally occurring variation in the number of integrated Maviruses. This allowed us to use the same genetic host background in all treatments, thus eliminating differences in the genetic background of the host and Mavirus populations. We tracked host, virus, and virophage population dynamics over 30 days in replicate chemostat cultures (approximately 150 host generations) under three different stressor levels that resulted in a gradient of integrated Mavirus per host cell. To look at the evolutionary consequences of differences in the frequency of integrated virophage and co-infection, we compared virus and virophage traits (replication and exploitation of virus by virophage) of ancestral (used to inoculate the chemostats) and evolved virus and virophage (from the end of the experiment).

## Materials and Methods

The experimental system consisted of the host *Cafeteria burkhardae* (strain E4-10P, hereafter: host) and the giant virus Cafeteria roenbergensis virus (CroV, hereafter: virus) and the virophage Mavirus (hereafter: virophage). All experiments were conducted using f/2 enriched artificial seawater medium (Guillard and Ryther 1962) supplemented with 0.025% (w/v) yeast extract. The host strain strain E4-10P used to start the experiments carried endogenous virophages (Hackl et al. 2021), but these did not produce particles under the experimental conditions of this study as has not been observed in similar experimental conditions (del Arco et al. 2024).

### Manipulation of frequency of integrated virophage

We used the antiviral oseltamivir as an off-target antiviral to manipulate the frequency of the integrated virophage. The mode of action of oseltamivir is described as the inhibition of neuraminidase enzyme which is involved in influenza A virus replication. While the influenza virus and the viruses used here are fundamentally different (e.g., RNA vs. DNA viruses), we found that the presence of oseltamivir at a predicted no effect concentration of 0.1mg/L led to significantly higher integration of the virophage into the host genome while showing no short effects on densities of host, virus and virophage (see Supplementary Information, Fig. S1 and Fig. S2). Note, that it is out of the focus of this study to determine the molecular mechanism of how oseltamivir affects Mavirus and CroV.

### Chemostat experiments

We assembled host-virus-virophage communities in chemostat systems with 400 mL f/2 enriched artificial seawater medium (Guillard and Ryther 1962) supplemented with 0.025% (w/v) yeast extract with a continuous flow through of 120 mL medium per day (= 0.3 dilution rate per day) (del Arco et al. 2020). Chemostats ran for 50 days in a culture room at 18 ± 0.5 ºC. Chemostats were inoculated with 7*10_3_ host cells/mL and isogenic virus and virophage were added after the host population dynamics had stabilized at days 20. We inoculated the viruses again on day 24 to ensure that the viruses established populations in all chemostats. Viruses and virophage were inoculated at a host:virus and host:virophage ratio of 0.1. The chemostats differed in the addition of the stressor to manipulate the integration of the virophage into the host genome. Four replicate chemostats received a one-time application of the stressor on day 26 (hereafter: *pulse* treatment). Four replicate chemostats received a constant inflow of the stressor with the inflow of medium at rate of 0.3 per day (hereafter: *disturbance* treatment). Four chemostat served as controls without the addition of the stressor. Analytical grade stressor (Sigma Aldrich, oseltamivir phosphate ≥98% HPLC) was diluted in f/2 enriched artificial seawater medium. The control chemostats received 2 mL of SW media at day 26.

We sampled chemostats daily (sample = 5 mL, except 10 mL once a week) for quantification of densities. Host densities were quantified from life samples using a hemacytometer and light microscopy. Virus and virophage samples were frozen for later DNA extraction (DNeasy 96 Blood & Tissue Kit, Qiagen, Hilden, Germany) and quantification by digital ddPCR (del Arco et al. 2023, 2024). All ddPCR results were analyzed using QUANTASOFT 1.7.4 The detailed methods and quality requirements for the data are described in the reference (del Arco et al. 2022).

Stressor concentrations were measured in randomly selected replicates on two days: day 26 right after the stressor pulse and the first addition to the disturbance treatment (4 chemostats receiving stressor pulse and one control) and day 29 as part of the daily sampling (in 2 replicates of the pulse and 3 replicates of disturbance treatment). Samples were kept at 4ºC and sent to SGS INSTITUT FRESENIUS GmbH (Radolfzell, Germany) to estimate concentration using gas Liquid Chromatography-Tandem Mass Spectrometry (LC-MS/MS). There was 0 mg/L in control chemostats, 0.11 ± 0.03 mg/L in the pulse and disturbance treatments after application (target concentration 0.1 mg/L). Six days after the first application, there was 0.05 ± 0.01 mg/L in the pulse and 0.16 ± 0.02 mg/L disturbance treatment.

### Integration of virophage into host genome

We measured virophage integration at day 22 (before addition of viruses) and by the end of the chemostat experiment at day 50. Virophage can be present in the community as free particles (horizontal transmission), or particle associated which can be either integrated virophage or virophage that is attached to organic particles. As the fraction of virophage attached to organic particles was negligible in our experiments (see Supplementary Information, Fig. S3), we used the particle-associated virophage fraction to estimate the average number integrated virophage per host (i.e., DNA copies per host). Specifically, virophage integration was estimated as the difference between total virophage abundance and virophage abundance after filtration through a 0.2 μm filter (Spartan® cellulose filter; Whatman®), which keeps the particles but let free virophage particles pass (Marvirus size ~70 nm (Fischer and Suttle 2011); for details see del Arco et al. 2022).

To test whether host survival in the presence of virus was influenced by the level of virophage integration, we compared host survival between ancestral and selected host population by the end of the experiment. We isolated 10 clones per chemostat from the last chemostat sampling day (day 50). We used serial dilutions from the treated samples for isolation of host clonal lines. Specifically, samples were diluted to 300 cell/mL, and 1 μL of the diluted sample was pipetted into 200 μL medium in 96 well plates at 18 ± 0.5 ºC. Growth of cells was checked after 6 days, and positive samples transferred to in 20 mL of SW media in tissue culture flasks to establish cultures of clonal lineages. We prepared isolated clones of host ancestor following the same protocol. We tested host survival by infecting the clonal cultures of the host (starting density 7*10_3_ host cells/mL) with virus at a host:virus ratio of 100 (n=2). Infections were done in 96 well plates (total volume of 200 μL of SW media). After 5 days post infection, host samples were fixed with Lugol’s solution (4% final concentration). Host survival was estimated using a high content microscope (ImageXpress Micro 4®; Molecular Devices, 20x magnification, transmitted light) and we applied a custom module with the MetaXpress® software for cell counting. We used presence/absence of host cells as measure of survival.

### Virophage evolution

We tested whether and how virophage evolved during the chemostat experiment by comparing traits (virophage replication and virophage inhibition of the virus (hereafter: virophage exploitation)) of ancestral virophages and virophages re-isolated from the end of the experiment. For the latter, we isolated virophages from the chemostats at day 50 by filtration through 0.2 μm filters. We then infected ancestral host in 200 μL of SW media in well plates with ancestral virus, and ancestral or re-isolated virophage at a host:virus and host:virophage ratio of 0.5 and 0 (as control). All combinations were replicated four times and started with a host density of 10_3_ cells/mL. After 3 days post infection we quantified virus and virophage densities (as DNA copies/mL) using ddPCR (see above). We assessed the replication and exploitation virophage traits between ancestral and selected virophages by comparing differences in DNA copy numbers after growing under standardized conditions. This allowed to identify heritable phenotypic changes in the virophage populations using standard experimental evolution procedures (Gómez and Buckling 2013; del Arco et al. 2024).

### Reactivation of integrated virophage

In addition, we isolated a total of 88 host clonal lineages from the end of the experiment: 23 came from the control, 28 from the pulse treatment and 37 from the disturbance treatment. We used serial dilutions from the treated samples for isolation of host clonal lines. Specifically, samples were diluted to 300 cell/mL, and 1 μL of the diluted sample was pipetted into 200 μL medium in 96 well plates at 18 ± 0.5 ºC. Growth of cells was checked after 6 days, and positive samples transferred to in 20 mL of SW media in tissue culture flasks to establish cultures of clonal lineages. As some cultures of the clonal lineages contained free virophage, we only continued with those cultures, where did not detect free virophage. We assessed virophage abundance following viral infection in ancestral and selected hosts. In replicated experiments (n=3), we either added ancestral virus (virus:host ratio of 20) or left hosts without virus. At 5 days post infection, we quantified virophage DNA copies using ddPCR (see above), and differences in virophage abundance were estimated as reactivated virophage by comparing DNA copies from the virus-free controls and the treatment where we added virus.

### Data analysis

All data analyses were performed in Rstudio (RStudio Team 2020) and R (R Core Team 2020) using the packages geepack (Højsgaard et al. 2020) and multcomp (Hothorn et al. 2008). Differences between models were considered relevant when p<0.05. We tested for differences in virophage integration by using generalized linear models (GLMs) (family = poisson) by treatment (control, pulse and disturbance) for day 50. To test for differences in host survival, we compared the percentage of clones growing in each treatment in the presence of ancestral virus using generalized linear models (family=poisson) using treatment (control, pulse, and disturbance) as explanatory variable. We evaluated differences in virophage reactivation between the ancestor and selected host clones with integrated virophages. We used generalized linear mixed model family=poisson) to compare between host lines (ancestral, control, pulse, and disturbance) and their interaction as explanatory variable and followed by a post-hoc tests including corrections for multiple testing (Tuckey). To test for host, virus and virophage population amplitude and density over the experiment, we used GLMs (family = poisson) with treatment (control, pulse, and disturbance) as explanatory variable. To test for differences in the ratios of virus:host, virophage:host and virus:virophage densities we used GLMs (family = poisson) with treatment (control, pulse, and disturbance) and time and their interaction as explanatory variable. We used wavelet analyses using the WaveletComp package (Roesch and Schmidbauer 2018) to estimate cycles length in population ratios of (a) virus-host, (b) virophage-host and (c) virophage-virus for each community. For this, data were smoothed following standard analysis procedures. Cycle lengths were compared between treatments (control, pulse, or disturbance) using linear models. All posthoc tests (Tuckey) corrected for multiple testing. Finally, to test for virophage trait evolution (virophage replication and exploitation of the virus), we used GLMs (family = poisson) with virophage line (ancestor, coming from control, pulse, or disturbance treatment) as explanatory factor.

## Results

To test for the relative role of the two modes of virophage replication, we manipulated the level of integrated virophage by adding a stressor and tracking the population dynamics of host, virus, and virophage over 50 days (30 days after virus and virophage inoculation to host populations) and compared the traits between the ancestral and re-isolated virophage. To test whether the manipulation was successful, we estimated the number of integrated virophages per host on day 22 immediately after virophage and virus inoculation and before stressor manipulation, as well as on day 50 at the end of the chemostat experiment. We found no integrated virophages on day 22. At day 50, virophage was integrated into the host genome in all communities and the integration was significantly higher in communities with the higher stressor manipulation (Fig. 1A, GLM, treatment, df=2, χ2 =-5226, p<2.2*10_−16_, post-hoc test, control vs. pulse vs. perturbation, p<2.2*10_−16_).

**Fig. 1.**
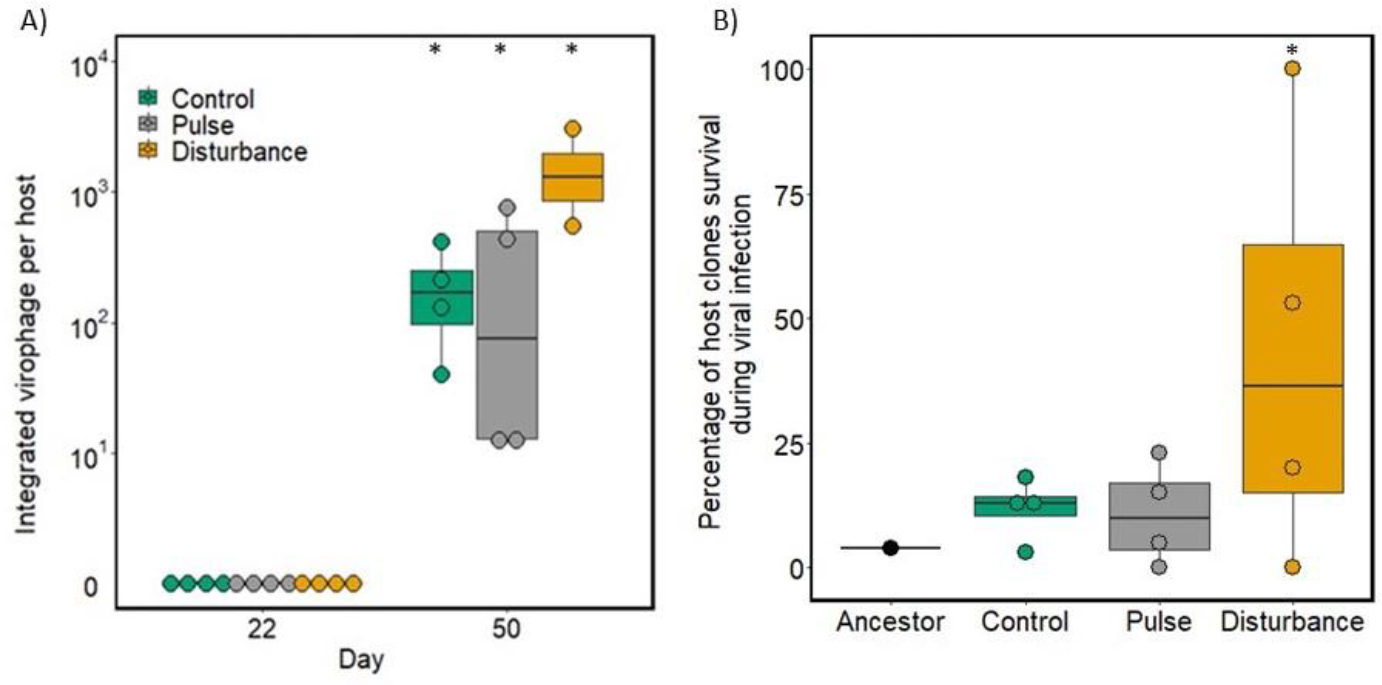
Virophage integration and host survival. A) Virophage integration as copies of virophage DNA per host genome after Mavirus and CroV addition into the chemostat communities at day 22 and at the end of the experiment at day 50. B) Host survival as the percentage of clonal lineages of the host in the presence of the ancestral virus. Asterisks above bars indicate statistical difference between treatments as determined by Tukey post hoc tests (p < 0.05).

We tested whether host survival following virus infection was affected by virophage integration using ancestral host clones (without integrated virophages) and selected host clones from day 50 (with integrated virophages via the chemostat experiment). The ancestral host clones did not survive viral infections, whereas selected host clones from the chemostats did. Specifically, selected host clones from communities with continuous antiviral exposure and higher virophage integration (perturbation) showed higher host survival than those from the pulse and control treatments (Fig. 1B, GLM: host line: df=3, χ2= 136.41, p<2.2*10-16; post-hoc test, perturbation differs from all others, p<2.2*10-16).

We assessed virophage reactivation in a subset of host clones selected from a collection of 88 clones (see Supplementary Information, Table S1). From this collection, we used host clones with integrated virophage and no signal of free virophage (4, 15 and 5 selected host clones from control, pulse, and perturbation communities, respectively) and ancestral host. We did not detect virophage reactivation in the ancestral host, whereas virophage reactivation in selected host clones was lower as the level of stressor increased from control to perturbation treatments (Supplementary Information, Fig. S4, generalised linear mixed model (GLMER), df=3, χ2= 38707731, p<2.2*10-16, post-hoc test, all comparisons, p<2.2*10-16).

Population densities cycled after virus and virophage introduction independent of the presence of the stressor (Figs. 3, S5). Cycles differed in their amplitude (Fig. 2A), with significant lower amplitudes in the host populations in the presence of the stressor (GLM: treatment: host, df=2, X_2_= 69881, p<2.2*10_−16_). Amplitudes of the virus population were also lower and significantly different in the present of the stressor (GLM: treatment: host, df=2, X_2_=411946915, p<2.2*10_−16_). For virophages, we found significantly higher amplitudes in the disturbance treatment (GLM: treatment: host, df=2, X_2_=1.8*10_10_, p<2.2*10_−16_). This resulted also in differences in the harmonic means of host, virus and virophage densities which differed between treatments (Fig. 2B, GLM: treatment: host, df=2, X_2_= 6380, p<2.2*10_−16_; virus, df=2, X_2_= 321638, p<2.2*10_−16_; virophage, df=2, X_2_= 3057765, p<2.2*10_−16_) with lower densities in the stressor treatments (post-hoc test: stressor manipulation: p<2.2*10_−16_).

**Fig. 2.**
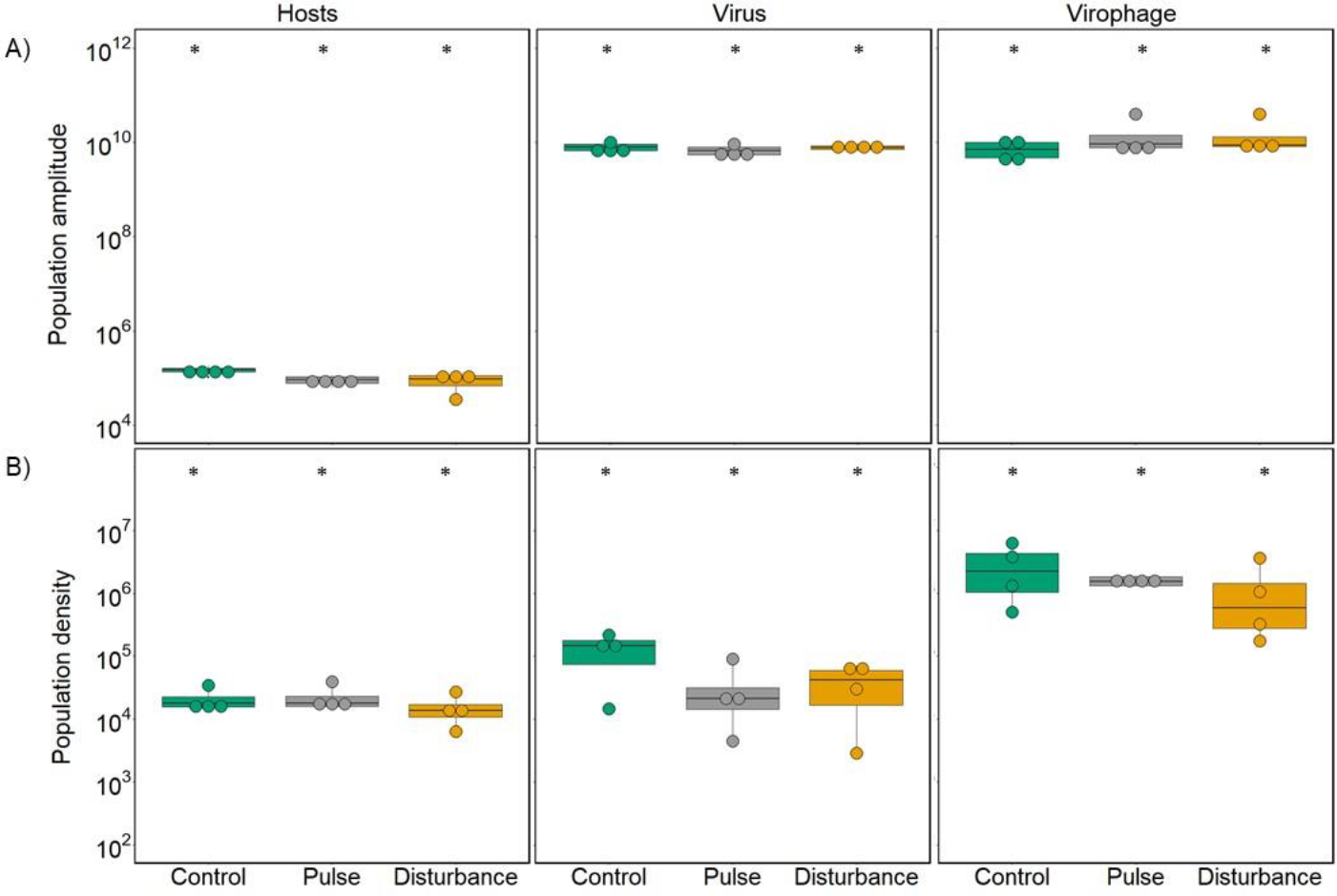
Host, virus and virophage amplitude and density. A) Amplitudes of population cycles (host cell/mL, virus or virophage DNA copies/mL over days 26-50 of the chemostat experiment (i.e., after the beginning of the stressor treatments). B) Harmonic mean densities (host cell/mL, virus or virophage DNA copies/mL over days 26-50 of the chemostat experiment (i.e., after the beginning of the stressor treatments). Asterisks above bars indicate statistical differences between treatments as determined by Tukey post hoc tests (p < 0.05).

**Fig. 3.**
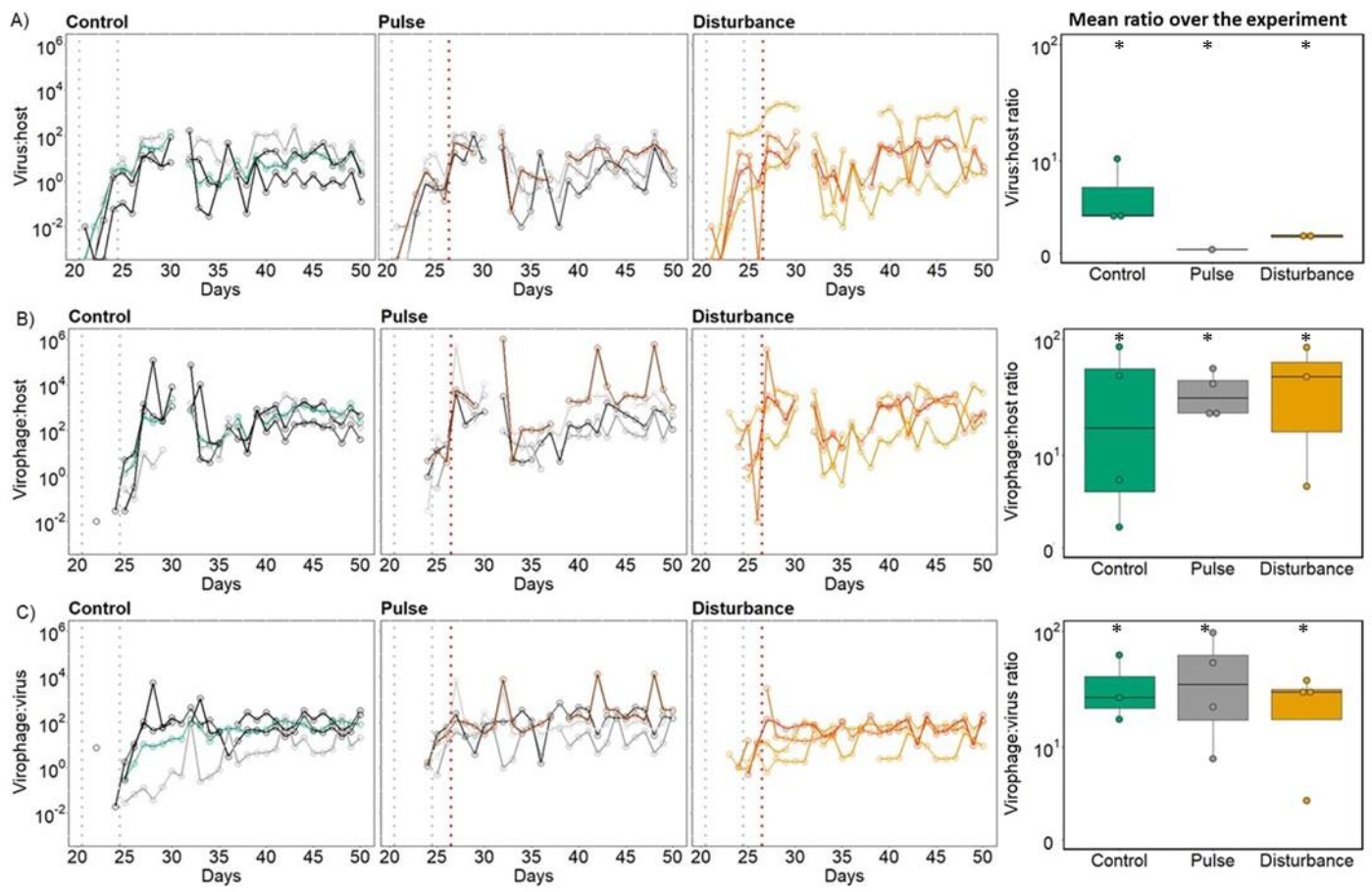
Population dynamics. Ratios of A) virus:host, B) virophage:host and C) virophage:virus in the control treatment stressor manipulation (first column), in the pulse treatment receiving a one-time stressor addition on day 26 (second column; red dotted line), and disturbance treatment where the stressor was added continuously after day 26 (third column). The 4 replicates per treatment and ratio are presented by different shades (green: control; grey: pulse, orange: disturbance). The grey lines represent virus and virophage addition. The last column of graphs represents the ratio harmonic mean for the 50 days of experiment in each treatment (control, pulse and disturbance).

Population ratios also differed between the communities, and they cycled over time with a significant period of 6 ± 1 days (Fig. 3) with no differences in cycle length between the treatments (virus:host, LM: treatment, χ_2_= 0.742, df=2, p = 0.599; virophage:host, LM: treatment, χ_2_= 1, df=2, p = 0.405; virophage:virus, χ_2_= 0.001, df=2, p = 0.999). The virus:host ratios changed over time depending on the treatment with lower average ratios in the presence of the stressor (GLM: treatment x time: df=2, X_2_= 15.943, p<0.003, treatment: df=4, X_2_= 21039, p<2.2*10_−16_, time: df=3, X_2_= 90.424, p<2.2*10_−16_; post-hoc test: all comparison: p=0.0001). The virophage:host ratio differed also over time and depending on the treatment with higher average ratios in the presence of the stressor (GLM: treatment x time: df=2, X_2_=269603, p<2.2*10_−16_, treatment: df=4, X_2_= 3935843, p<2.2*10_−16_, time: df=3, X_2_= 656096, p<2.2*10_−16_; post-hoc test: all comparison: p<2.2*10_−16_). This also led to significant differences for the virophage:virus ratios over time and between treatments with significant higher ratios in the pulse treatment (GLM, treatment x time: df=2, X_2_= 5758.8, p<2.2*10_−16_, treatment: df=4, X_2_= 50442, p<2.2*10_−16_, time: df=3, X_2_= 6773.4, p<2.2*10_−16_; post-hoc test: all comparison: p<2.2*10_−16_)

We tested whether and how virophage reproduction and exploitation evolved in the experiment by comparing these traits for virophages re-isolated from day 50 of the experiment to the virophage used to start the experiment. Reproduction of re-isolated virophages was higher compared to ancestral virophages and differed among the treatments, with decreasing reproduction with increasing stress level (Fig. 4A, GLM: virophage line: χ_2_ = -46743054, df=3, p < 2.2*10_-16_, post-hoc test: all comparison: p<2.2*10_−16_). Virophage exploitation of the virus was lower for re-isolated virophages compared to ancestral virophages (Fig. 4B, GLM: virophage line: χ_2_ = -10902981, df=3, p < 2.2*10_-16_; post-hoc test: all comparison: p<2.2*10_−16_) and lower despite of higher virophage replication of virophages in selected virophages (Fig. 4A).

**Fig. 4.**
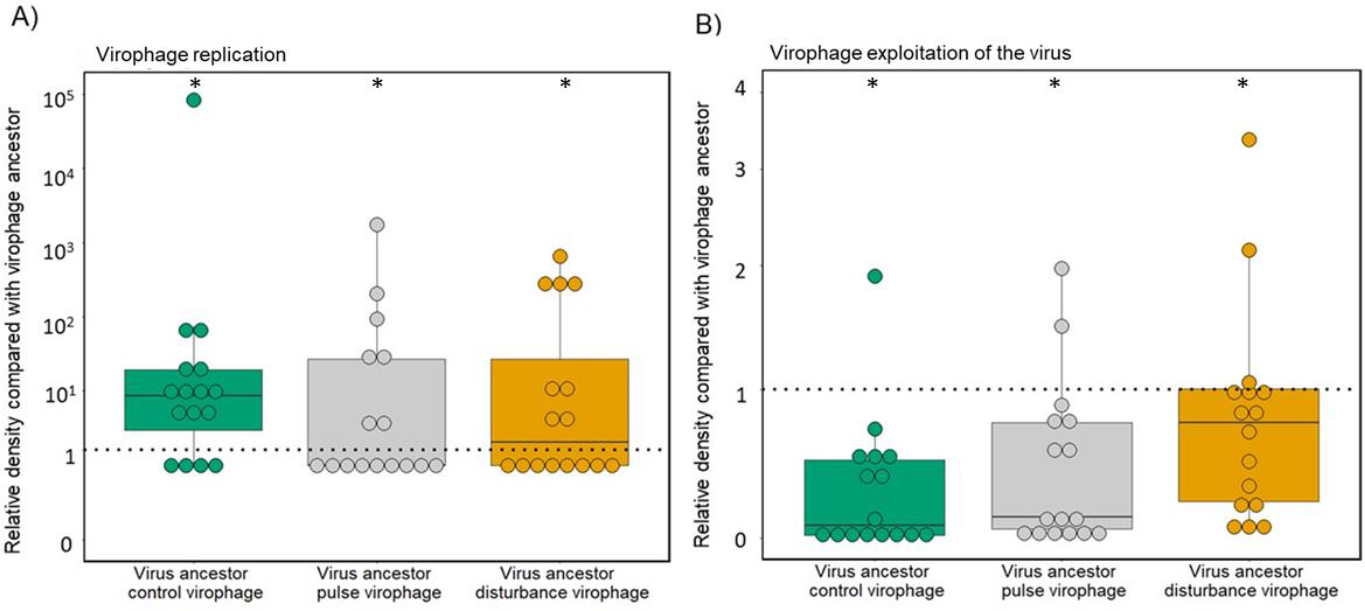
Virophage evolution. Differences in virophage replication and virophage exploitation of the virus of selected virophage respect to ancestral virophages. A) Virophage replication estimated as virophage relative densities of selected virophage 3 days post infection respect to ancestral virophage density in infections with ancestral host and virus. B) Virophage exploitation estimated as its effect on virus replication comparing ancestral or selected virophages. A value below 1 means that the re-isolated virophages have increased virus exploitation compared to the ancestor. In both panels green represent virophage isolated from control treatment; grey, virophage isolated from pulse treatments; and orange, virophage isolated from disturbance treatments). Each point represents an isolated virophage population coming from one of the 4 replicates per chemostat (trait assays with 4 replicates; total 16 virophages per treatment). Asterisks above bars indicate statistical differences between treatments as determined by Tukey post hoc tests (p < 0.05).

## Discussion

The dual nature of the virophage lifestyle is unique among eukaryotic DNA viruses and provides the virophage with intricate ways to interact with the virus and the host cell with distinct effects on the virophage and virus reproduction. The relative roles of the different replication modes can change with conditions, and their shifts may play a central role in the ecological and evolutionary dynamics of the community because the mode of virophage replication affects virus replication. Here, we tested the ecological and evolutionary consequences of different virophage replication modes by indirectly manipulating the frequency of integrated virophages per host cell.

We observed that virophages evolved higher levels of replication but lower levels of exploitation over time in all treatments, as also previously observed in this system in the absence of a stressor (del Arco et al. 2024). Compared to the ancestor the increase in replication and decrease in exploitation of the virus was less pronounced at higher levels of virophage integration. This is consistent with the prediction for the effect of higher levels of integrated virophage on virophage evolution. Low levels of integrated virophage in the absence of the stressor impose stronger selection on the virophage for higher replication, as horizontal transmission would be the alternative mode to persist over vertical transmission. Increased virophage replication, which promotes coinfections and subsequently inhibits viruses, may ultimately result in virus extinction and a decline in virophage populations. This scenario promotes evolution towards lower virophage exploitation of the virus as observed. In the presence of the stressors, the high levels of integrated virophage mitigate selection pressure on virophage exploitation trait (being more similar to ancestral virophages compared to the changes in populations with low levels of integrated virophage). Reactivation of integrated virophages, which facilitates replication for both viruses and virophages, allows virophage replication through coinfections or new integrations alleviating selection pressures on virophage exploitation.

The pattern of increased replication and decreased exploitation shows a correlation between exploitation of the virus and virophage replication, where virophage with higher replication have lower virus exploitation. Trait correlations are often observed in microbial systems including viruses with strong effects on the evolutionary and ecological dynamics (Friman and Buckling 2013; Koskella and Brockhurst 2014). For example, they can influence the rate of evolution of resistance mechanisms against multiple consumers (Ford et al. 2016), the direction of evolutionary change (Scheuerl et al. 2019), and they can determine population dynamics directly and indirectly through eco-evolutionary dynamics (Frickel et al. 2017). Here the virophage fitness is expected to be optimized when the virophage can parasitize the virus, instead of only vertical transmission with host replication. High levels of exploitation would lead to very low virus densities or even extinction over time (Barreat and Katzourakis 2024), reducing virophage fitness. Experimental and field data suggest that virophages are key players shaping the dynamics and evolutionary trajectories of host and virus populations (La Scola et al. 2008; Yau et al. 2016; Fischer and Hackl 2016; del Arco et al. 2022). For instance, in natural communities, virophages that infect algae can exacerbate blooms of Antarctic phytoplankton during summer periods by suppressing parasitic viruses (Yau et al. 2016). Virophages of heterotrophic flagellates, which play a key role as bacterivores, have the potential to impact biogeochemical cycles and disrupt the regulation of flagellates by their viruses. (Massana et al. 2007).

Host, virus and virophage populations showed significant and similar cycles over time across communities despite of virophage trait differences depending on stressor manipulation. This suggests diverse underlying mechanisms driven by trait variation resulting in similar population dynamics. Most studies modelling host-virus-virophage interactions find cyclic dynamics in their simulations (Wodarz 2013; Taylor et al. 2014; Barreat and Katzourakis 2024) that can stabilize for example depending on the virophages inhibition of the virus (Wodarz 2013; Barreat and Katzourakis 2024), and co-infection mode (together vs. sequential Taylor et al. 2014). Reduced exploitation in consumer-resource systems can stabilize the dynamics of consumer-resource systems, leading to smaller amplitudes or a shift towards steady-state dynamics (Rosenzweig and MacArthur 1963) and a recent modelling study showed that increasing the inhibition of virophages on virus replication can have a stabilizing effect on the oscillatory dynamics (Barreat and Katzourakis 2024). Here we did not find a pattern suggesting stabilization due to differences in virophage exploitation of the virus. Instead, all three populations cycled but differed in their amplitudes dependent of treatment, rather than having smaller amplitudes with higher levels of exploitation. This observation could be explained by at least three non-exclusive mechanisms, that needs further studies. First, the trait differences we observe at day 50 of the experiment might only reflect very recent evolutionary changes, and differences in population dynamics can only be observed over longer experimental time scales. Second, the relative roles of virophage reproduction modes – vertical transmission, coinfection or reactivation of integrated virophage-changed over time as a function of stressor treatments and population dynamics. However, the consequences for population dynamics have not been fully explored theoretically. Finally, stabilization with reduced exploitation may be less common in systems with such complex interactions as that of the virophage with the virus and the host, and the parameter space explored in the models does not cover some of the key biological parameters. Models capable of fitting experimental population trajectories could help to identify the underlying mechanisms governing the structure and function of host-virus-virophage communities. For example, they could help predict which population dynamics are indicative of the community transitioning from a region of coexistence to extinction as a function of infection mode.

The ddPCR protocol used in this study targets the virophage Mavirus, but its efficiency to detect other endogenous Mavirus-like elements (EMALEs) carried by C. burkhardae strain E4-10P (Hackl et al. 2021) has not been assessed. It is thus possible that other EMALEs have been reactivated affecting the ecological and/or evolutionary dynamics. Koslová et al. (2024) characterized the reactivation of EMALES in a collection of globally distributed Cafeteria populations and they found that reactivation was stochastic, inefficient and EMALE-virus specific (e.g., only one out of eight EMALEs reactivate upon CroV infection). For the host strain E4-10, which we used here, EMALE04 was reactivated by CroV infections but low copies were produced, and host died within several days after infection. However, they also showed that EMALE04 inhibition of the virus can amplify in subsequent rounds of infection reaching similar inhibition levels of the virus population as Mavirus (Koslová et al. 2024). Whether and how EMALE04 reactivated and competed with Mavirus in our experiment is not known. Since EMALE04 reactivation has been shown to be stochastic, we would expect that EMALE04 reactivation and accumulation in the communities would lead to large variation across replicates within treatments, which we don’t observe. Competition among virophages is unexplored, which is particularly compelling due to its potential implications for understanding the eco-evolutionary dynamics of host-viral-virophage populations.

Virophage integration levels are influenced by environmental conditions, as demonstrated by our stressor manipulation. However, there are other factors that can influence integration levels and thus their role in protecting the host population. These include host cell cycle and physiological states, or the genetic background of the host, virus and virophage. We found that higher level of integration showed a positive correlation with higher host survival, negative correlation with virophage reactivation and with viral inhibition. As host-virus-virophage communities are characterised by a high degree of population interdependence, shifts in the relative roles of replication modes must be considered as a potential key mechanism influencing community ecological and evolutionary responses.

## Acknowledgements

We thank Matthias Fischer and his Lab for helpful comments on the experimental data.

## Author contribution

AdA, LB, conceived the study, AdA, LB designed of the study; AdA carried out the experiment, AdA, LB analysed the data; AdA, LB wrote the manuscript, all authors edited the manuscript.

