## Supplementary Information for "Virophage replication mode determines ecological and evolutionary changes in a host-virus-virophage system"

***Effect of oseltamivir on virophage integration***

To test for the effect of oseltamivir on virophage integration, we estimated the number of integrated virophages per host cell in short term experiments where we exposed hosts (starting density: $10^5$ cells/mL) to two concentrations of oseltamivir (0 and 1.0 ug/L) in the presence (host:virophage ratio of 10) and absence of virophage (host:virophage ratio of 0) for for 8 days (~32 host generations). Experiments were done in 10 mL of SW media in tissue culture flasks at 18 ± 0.5 ºC. After 8 days of culturing, which is sufficient time to observe virophage integration into the host genome (Fischer and Hackl 2016), we sampled two fractions from the cultures for virus quantification by ddPCR (see main text). First, we extracted DNA from the whole sample, representing the total virophage population (i.e., free virions and particle associated viruses). Second, we extracted DNA from the filtrate after filtering through 0.2 µm filter representing the free virions. The particle associated fraction is then calculated from the difference in DNA copies/mL between the two fractions. Host samples were fixated with Lugol's solution (4% final concentration) and quantified later using a hemacytometer and light microscope (20x magnification). Using the information on the DNA copies/mL and host cells/mL we estimated the average number of integrated virophages per host cell. We found integrated only in the presence of the virophage (Fig. S1, Generalized Linear Model (GLM), treatment x stressor: df=1, $X^2$= 9.4902x$10^{-8}$, = 0.9998, treatment: df=1, $X^2$= -14.778, p<0.001) with significantly higher integrated virophage per host in the presence of the stressor (Fig. S1, GLM, stressor: df=1, $X^2$= -37.43, p<0.001).

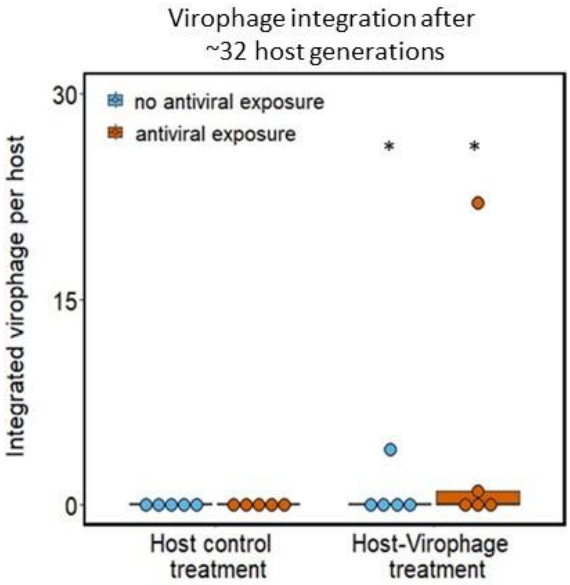


Fig. S1 **Manipulation of virophage integration.** Virophage integration as DNA copies of virophage per host after 8 days post-infection with Mavirus in the presence and absence of the stressor. Asterisks above bars indicate statistical differences between treatments as determined by Tukey post hoc tests ($p < 0.05$).

***Effect of oseltamivir on host, virus and virophage population growth***

We also tested whether exposure to oseltamivir influences the population growth of host, virus and virophage. For this we grew i) host alone (control), ii) host and virus, and iii) host, virus and virophage in 15 mL of SW media in tissue culture flasks at 18 ± 0.5 $^{o}$C for three days in replicates of four in the absence and presence of the stressor (0 and 1.0 ug/L of oseltamivir). All combinations were started with a host density of $10^5$ cell/mL and viruses and virophage were added at a host:virus:virophage ratio of 5. After 3 days, we sampled and quantified hosts, viruses and virophages as described in the main text. We found that population densities were not affected by the presence of the stressor after three days of growth, which represent approximately 1 round of replication of the virus and virophage (Fig. S2; Kruskal test: host, df=1, $X^2$=3.125, p=0.0771; virus, df=1, $X^2$=2, p=0.1573; virophage, df=1, $X^2$=2, p=0.1573).

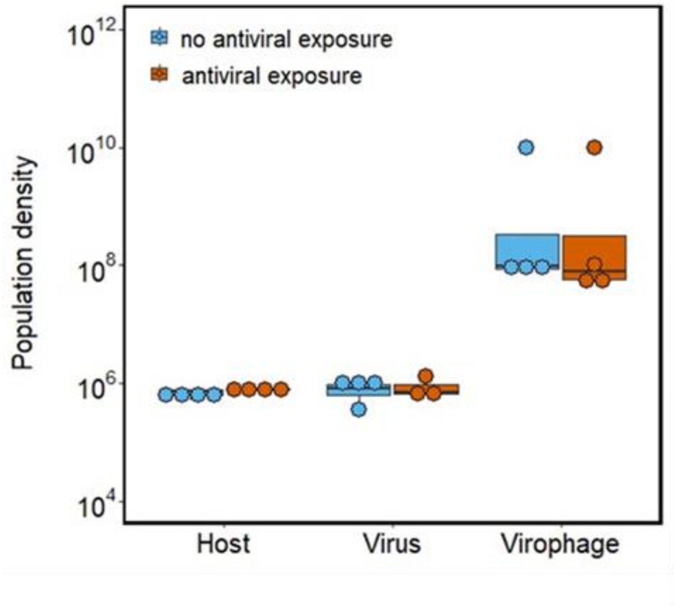


Fig. S2 **Host, virus and virophage densities in the absence and presence of the stressor**. Density of host (cells/mL) and virus (DNA copy numbers/mL) 3 days post-infection in the absence and presence of the stressor.

***Effect of oseltamivir on virus and virophage aggregation***

The particle associated fractions contains the DNA copies integrated into the genome of the host cells, and potentially virions attached to organic particles (e.g., bacteria or cell debris). We tested for the attachment to organic particles in a separate experiment without hosts. We exposed virus and virophage together in culture flasks (n=4) to two stress environments (0 and 1.0 ug/L of oseltamivir) for 24 hours under gentle shaking conditions (160 rpm). We followed the same protocol as above with the difference of using a filter with a pore size of 0.45 µm. We then used the information on DNA copies/mL in the filtrate and the total sample (without filtration) as a proxy for virions attached to organic particles (or being stuck in the filter). We found that the presence of the stressor did not result in density differences of free virus (Fig. S3, stressor, kruskal test: df=1, $X^2$= 0.035714, p = 0.8501) and virophage particles (Fig. S3, stressor, kruskal test: df=1, $X^2$= 0.34146, p = 0.559), suggesting that the presence of the stressor did not affect aggregation and attachment to organic particles. As we did not quantify the fraction of virions attached to organic particles or to hosts (including the integrated virophage), our estimate of virophage per host is likely an overestimation. There should be however no systematic differences between the stressor treatments, as we do not see an effect of the stressor (Fig. S3).

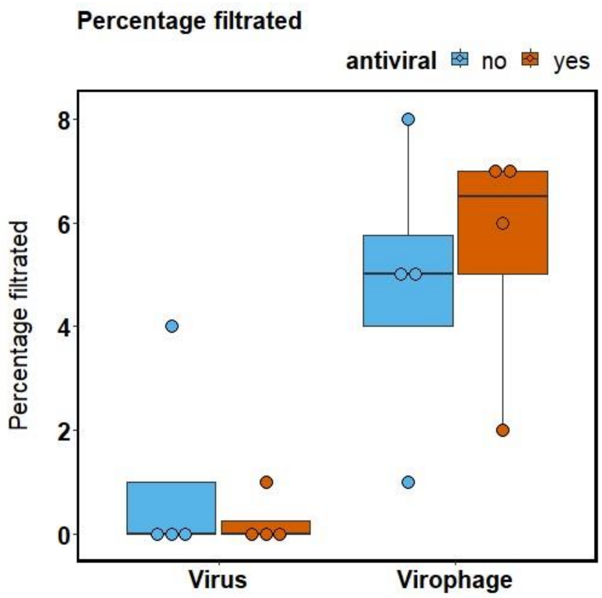


Fig. S3 **Virus and virophage aggregation to particles in the absence and presence of the stressor**. Virus and virophage free particles estimated as the percentage of particles going across a filter of 0.45 µm in the presence and absence of the stressor (n=4).

Table S1. Free virus, free virophage and integrated virophage average DNA copy number in the total 88 host selected clones (DNA copies/mL, mean ± se) from control (23 clones), pulse (28 clones) and disturbance (37 clones).

| Antiviral treatment | Free virus | Free virophage | Integrated virophage |
|---|---|---|---|
| **Control** | $8,2*10^{4} \pm 4,4*10^{4}$ | $1,8*10^{5} \pm 1,2*10^{5}$ | $1,4*10^{8} \pm 5,5*10^{7}$ |
| **Pulse** | $7,0*10^{4} \pm 2,7*10^{4}$ | $2,0*10^{5} \pm 1,8*10^{5}$ | $2,1*10^{8} \pm 5,9*10^{7}$ |
| **Disturbance** | $4,9*10^{4} \pm 1,3*10^{4}$ | $1,8*10^{5} \pm 3,8*10^{5}$ | $1,6*10^{8} \pm 4,7*10^{7}$ |

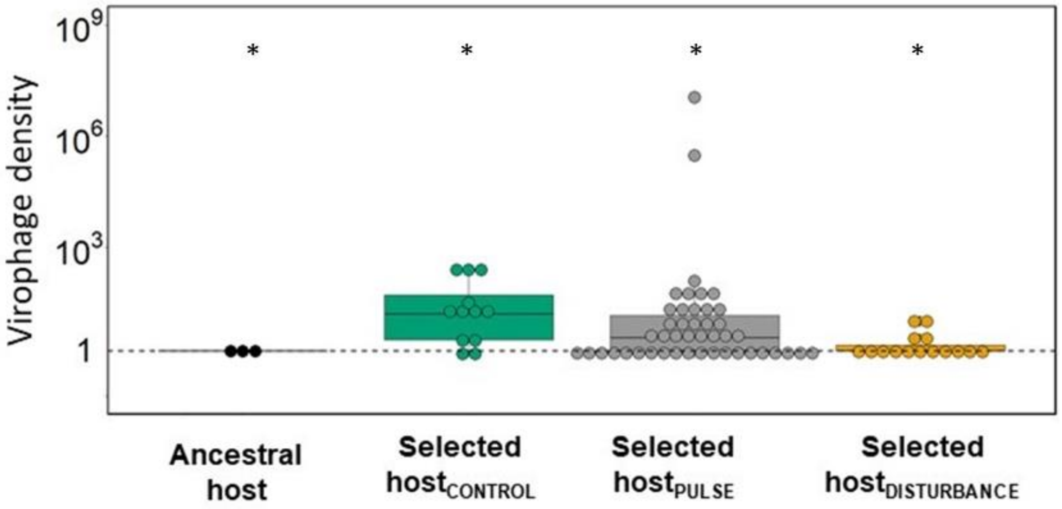


Fig. S4 **Virophage reactivation**. Differences in virophage density (DNA copies/mL) as a proxy of virophage reactivation. Virophage density after 5 days post infection in ancestral host or selected host clones in viral treatments compared to control treatments without viral infections. Black colour represents the ancestral virophage; green colour represents virophage isolated from control treatment; grey, virophage isolated from pulse treatments; and orange, virophage isolated from disturbance treatments. Each point represent one replicate (n=3) of virophage abundance coming from ancestral host clones or from selected host clones coming from the different chemostat communities varying in antiviral exposure. The central dotted line represents virophage density in ancestral host or selected host clones not infected with virus. Asterisks above bars indicate statistical differences between treatments as determined by Tukey post hoc tests ($p < 0.05$).

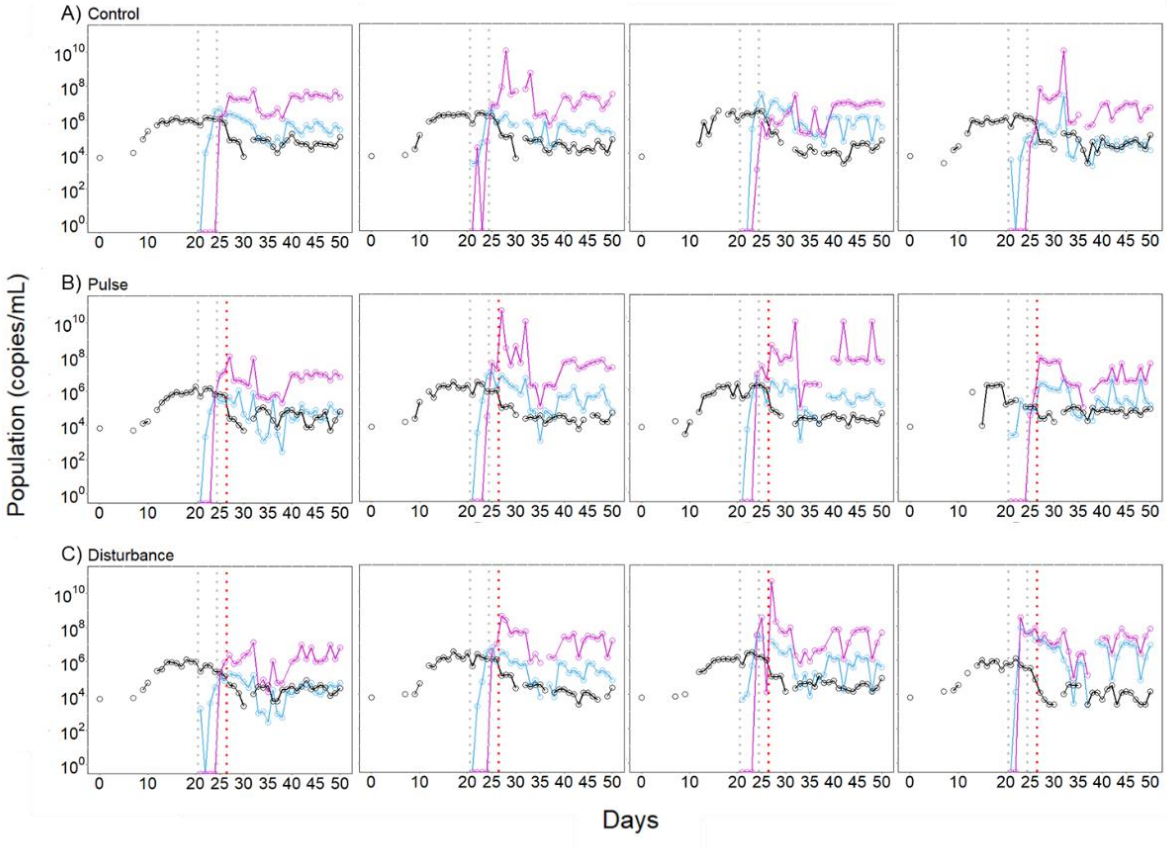


Fig. S5 **Population densities of host-virus-virophage over time.** Population densities A) in control treatment stressor manipulation (no antiviral addition), B) in the pulse treatment receiving a one-time stressor addition on day 26 (black dotted line), C) and disturbance treatment where the stressor was added continuously after day 26. Black, host; blue, virus; purple, virophage. Host was allowed to establish in the chemostats before virus and virophage were added on day 20 and 24 (grey vertical dotted lines).
